# Evaluation of Known Human PDE Inhibitors Against Nematode PDE4s

**DOI:** 10.1101/2023.02.24.529780

**Authors:** Clianta S Anindya, Charles S Hoffman

## Abstract

Parasitic nematodes are responsible for more than one and a half billion infections world-wide. The drugs developed against these infections only target a few different proteins. As drug resistance is becoming more common, there is a need to develop new drugs against new targets. Cyclic Nucleotide Phosphodiesterases (PDEs), are enzymes that hydrolyze the cyclic molecules of cyclic AMP and cyclic GMP. Physical properties of mammalian PDEs have led them to become well-established as drug targets. Mammals possess 11 families of PDEs, many of which are the target of selective and potent drugs. Nematodes have 6 PDE genes representing 6 families, which have not been well-studied; *C. elegans*, is a model organism nematode that would allow people to assess the therapeutic benefit of targeting PDEs. The Hoffman Lab has developed a platform for discovering PDE inhibitors and has carried out high-throughput screens (HTS) to help identify inhibitors of mammalian PDE4, PDE7, PDE8, and PDE11 families. The PDE4 family in *C. elegans* is of particular interest as work in *C. elegans* suggests that it may be involved in neuronal function. However, research has shown that two compounds developed against mammalian PDE4s generally do not work on *C. elegans* PDE4. Therefore, the goal of this project is to screen a collection of compounds discovered by the Hoffman Lab to identify the compounds that will affect *C. elegans* or parasitic nematode PDE4s to find compounds that could then be tested for their effect on *C. elegans* and parasitic nematodes. This research could then identify an effective new target for drug development to treat infections by parasitic nematodes.

**Summary:** Parasitic nematodes are the soil worms responsible for more than one and a half billion infections around the world. While drugs are being developed against them, these drugs are designed against relatively few proteins, which is a problem as drug resistance becomes more common. Therefore, there is a need for new drugs. PDEs are enzymes that hydrolyze the signaling molecules cAMP and cGMP. Mammalian PDEs have been well-established as drug targets. In nematodes, there are 6 PDE genes representing 6 families of the 11 families found in mammals. Additionally, a free-living model organism nematode, *Caenorhabditis elegans* (*C. elegans*) can be used to assess the impact of PDE inhibition on nematode biology. In the Hoffman Lab, they have developed a platform for discovering PDE inhibitors and have used these in high throughput screens to identify inhibitors of mammalian PDE4, PDE7, PDE8, and PDE11 families. The PDE4 family in *C. elegans* is of particular interest as work in *C. elegans* suggests that it may be involved in neuronal function. However, research has shown that compounds developed against mammalian PDE4s generally do not work on *C. elegans* PDE4. Therefore, by the end of this project we hope to identify the compounds that do work on nematode PDE4s that could be used to test whether they have the potential to treat these infections.

## 1 Introduction

Parasitic nematodes are the soil worms responsible for more than one and a half billion infections around the world. The existing drugs developed against these infections can only target a few different proteins and drug resistance becoming is becoming more common. According to the new World Health Organization (WHO) roadmap, new drugs or drug regimens that kill or permanently sterilize adult filarial worms would significantly improve elimination timelines, and accelerate the achievement of the program goal of disease elimination [1].

*C. elegans* is a free-living nematode, it has become an important model organism as it is an outstanding experimental system, due to its small size, rapid life cycle, transparency, and well-annotated genome, which can be useful in research laboratories [2]. Therefore, it can be used to evaluate potential anti-nematode drugs [3] [4].

Cyclic Nucleotide Phosphodiesterases (PDEs) are a class of related enzymes that catalyze the hydrolysis of the 3’ cyclic phosphate bond of adenosine 3’5 monophosphate (Figure 1). PDEs work by selectively catalyzing the hydrolysis of the 3’ cyclic phosphate bond of cAMP [5].

**Figure 1:**
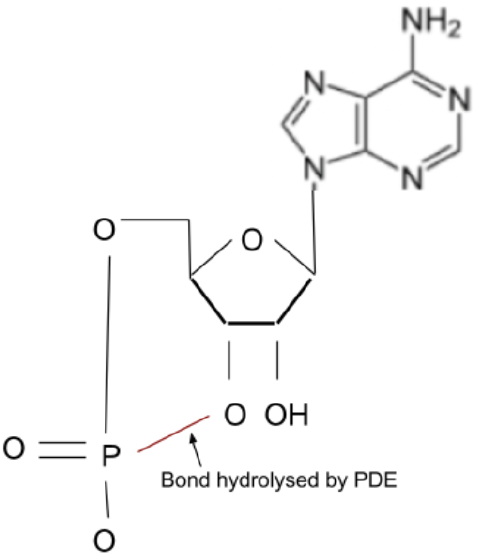
PDEs convert cAMP to AMP by catalyzing the hydrolysis of the 3’ cyclic phosphate bond.

Several different PDEs can be expressed by any single cell type and the principle inhibition of PDE activity could be a valid therapeutic target, because the regulation of the degradation of any ligand or second messenger can often make a bigger change in concentration than those of synthesis rates [5]. The huge clinical success of PDE inhibitors (especially PDE5 inhibitors) has created continued interest in PDE inhibitors as potential therapeutics and in overcoming the limitations of current PDE inhibitors [6]. Mammals possessed 11 families of PDEs, many of which are the target of selective and potent drugs. Nematodes have 6 PDE genes representing 6 families. In cyclic nucleotide signaling, it was found that *C. elegans* PDE4s might be important for neuronal health [7]. However *in vitro* enzyme assays of roflumilast and zardavarine, both of which are potent inhibitors of mammalian PDE4s, are 100-fold less effective on *C. elegans* PDE4 [8].

Chemical genetics, both *in vitro* biochemical and phenotypic assay platforms have clear limitations in high-throughput screening (HTS) for drug discovery. Yeast based screens against heterologously-expressed proteins have the potential to bridge the gap between phenotypic and biochemical assays for HTS [9]. This flexibility ensures that a wide variety of targets, including those regarded as less tractable (e.g., transmembrane proteins), can be screened in ultra-high-throughput screening (uHTS) in a robust and cost-effective manner [10].

The hypothesis of this project is that compounds designed against mammalian PDE4s do not necessarily work on nematode PDE4s. Therefore, the goal of this project is to screen a collection of compounds discovered by the Hoffman Lab to identify ones that will affect *C. elegans* or parasitic nematode PDE4s to find compounds that could then be tested for their effect on *C. elegans* and parasitic nematodes.

## 2 Materials and Methods

### 2.1 Strains

Strains used in this project are listed in Appendix (A.1).

### 2.2 Yeast Growth Media

Yeast were grown on YES, EMM, SC-ura or 5FOA medium as previously described [11] [12].

### 2.3 Compounds

Compounds used in this project are listed in Appendix (A.2). Compounds designated as BC are referred to ones identified in the high-throughput screening (HTS) by the Hoffman Lab.

### 2.4 SC-ura Halo Assay Test

In SC-ura halo assays, we are looking for a halo. Halo is a circle in which colony growth is reduced or eliminated, it is a sign of PDE inhibition that leads to a loss of growth in the area of the active compound due to repression of the *fbp1-ura4* reporter (Figure 2).

**Figure 2:**
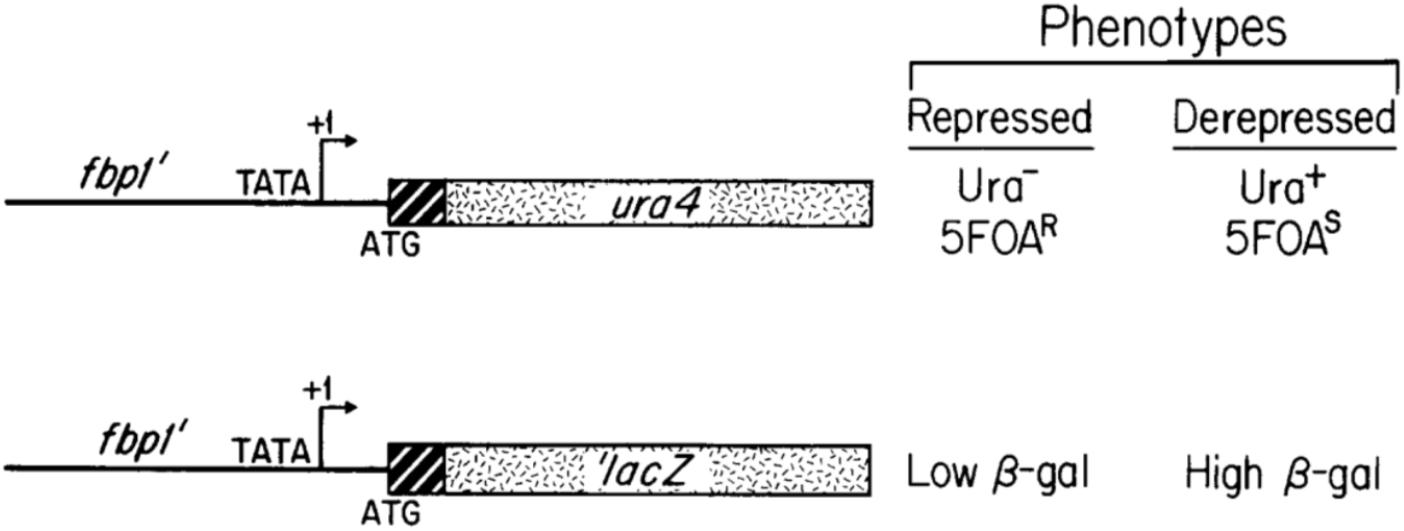
The structure of the *fbp1-ura4* and *fbp1-lacZ* fusions and their associated with phenotypes. The *fbp1-ura4* and *fbp1-lacZ* are both translational function which include the first four condons of the *fbp1* open reading frame. The *S. pombe ura4* gene encodes OMP decarbocylase which is required for uracil prototropy and for sensitivity to 5FOA; the *E. coli lacZ* gene encodes *β – galactosidase*[11].

Cells are grown in an EMM medium and approximately 10,000 cells are spread to an SC-ura plate. One microliter of a compound is then spotted onto the plate and plates are incubated at 30ºC for three or four days before photographing.

### 2.5 5FOA Assay Test

In 5FOA assay test, we are looking for the PDE inhibitors that could help the 5FOA assays from no growth to growth on a medium containing 5FOA. The 5FOA assay uses strains whose PKA activity is insufficient to repress expression of the *fbp1-ura4* reporter [11], leading to a loss of growth in 5FOA-containing medium (Figure 3A). Inhibition of the PDE elevates cAMP levels, leading to 5FOA-resistant growth (Figure 3B).

**Figure 3:**
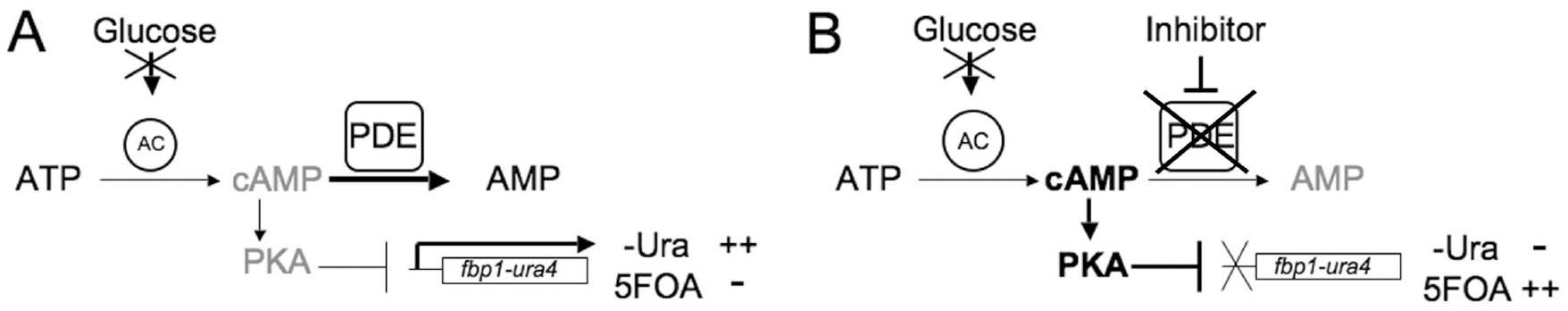
**(A)** Cells carrying mutations in genes required for glucose signaling have reduced adenylyl cyclase activity to lower cAMP levels. This results in low PKA activity and a failure to repress *fbp1-ura4* transcription. These cells grow in medium lacking uracil (–Ura) but do not grow in medium containing 5FOA. **(B)** A screen for PDE inhibitors can be carried out by taking a strain such as the one in panel A and screening for compounds that enhance growth in 5FOA medium [13].

Strains are grown to exponential phase in the EMM medium containing 10 mM cAMP to ensure the activation of PKA. Cells are collected and washed in 5FOA medium cell and then resuspended to a density of 2.5 × 10^4^ to 2 × 10^5^ cells per ml, depending on the strain. Next, 40 μl cultures are placed in a 384-well microtiter plate in the presence of compound ranging from 50μM to 1.6μM, along with a DMSO vehicle control. It is then grown for 48 hours at 30ºC before reading for optical density.

Upon testing 5FOA assays, we are also trying to adjust the 5FOA condition (Appendix A.4) in order to see the optimal condition to work with. The two conditions we adjusted were the amount of 5FOA and the cell density.

## 3 Results

### 3.1 PDE Inhibition using Halo Assay Results

Seventy-seven compounds were tested (Appendix A.2) for their ability to form halos using strains expressing mammalian PDE4B2 or PDE7A1 (data not shown). Of these, 40 compounds were selected for testing on strains expressing human PDE4B2, *C. elegans* PDE4, *C. elegans* PDE5, and *Ancylostoma ceylanicum* (Acey) PDE4 (Figure 4). While many compounds produced halos on the strain expressing human PDE4B2, only BC51 and BC65 produced halos on strains expressing the nematode PDEs.

**Figure 4:**
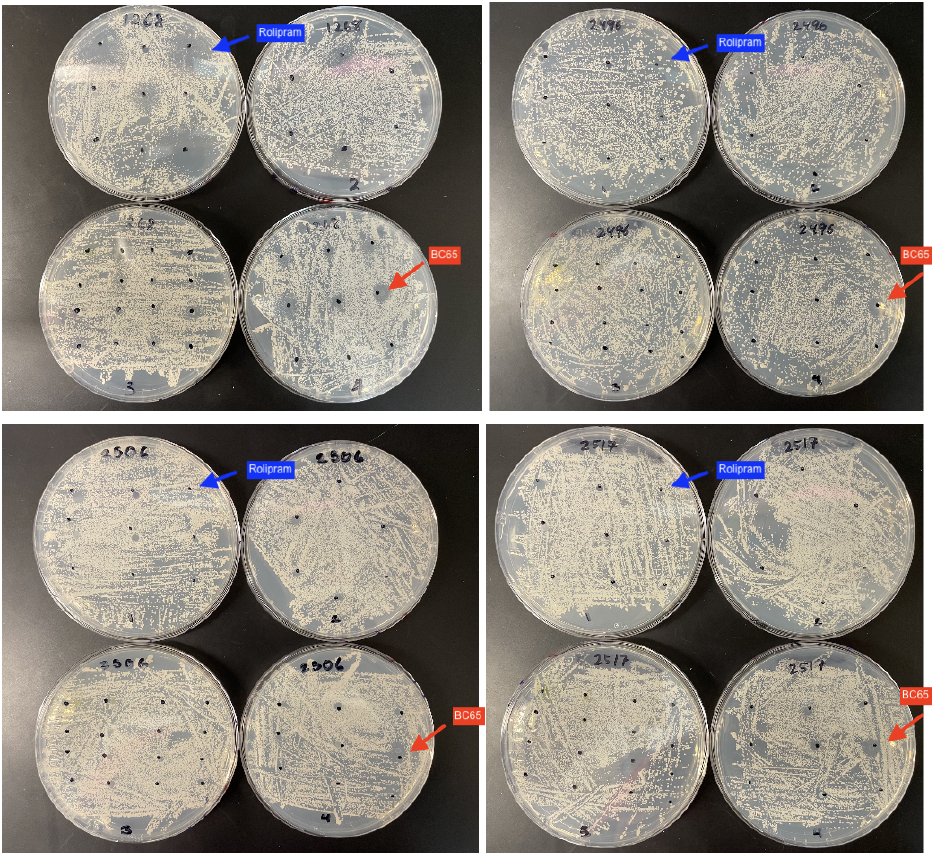
SC-ura halo assays reveal differential sensitivity to BC65 by human PDE4B22, *C. elegans* PDE4, *C. elegans* PDE5, and *Ancylostoma ceylanicum* (Acey) PDE4. Strains CHP1268 (human PDE4B2-upper left plate), CHP2496 (*C. elegans* PDE4-upper right plate), CHP2506 (*C. elegans* PDE5-lower left plate), and CHP2517 (*Ancylostoma ceylanicum* (Acey) PDE4-lower right plate) were plated to SC-ura plates. Plates received one microliter spots of test compounds and then incubated for three days at 30°C before photographing. PDE inhibition leads to a loss of growth in the area of the active compound. The red arrow denotes compound BC65 that shows halos on all four strains while the blue arrow denotes compound Rolipram that only shows a halo on the strain expressing human PDE4B2.

After the 40 evaluated compounds were tested, we then tested compounds Rolipram, BC51, and BC65 against human PDE4B2, *C. elegans* PDE4, *Ancylostoma ceylanicum* (Acey) PDE4, *Brugia malayi* (Bmal) PDE4, *Onchocerca volvulus* (Ovo) PDE4, *Strongyloides ratti* (Strat) PDE4, *Angiostrongylus costaricensis* (Acos) PDE4 (Figure 5).

**Figure 5:**
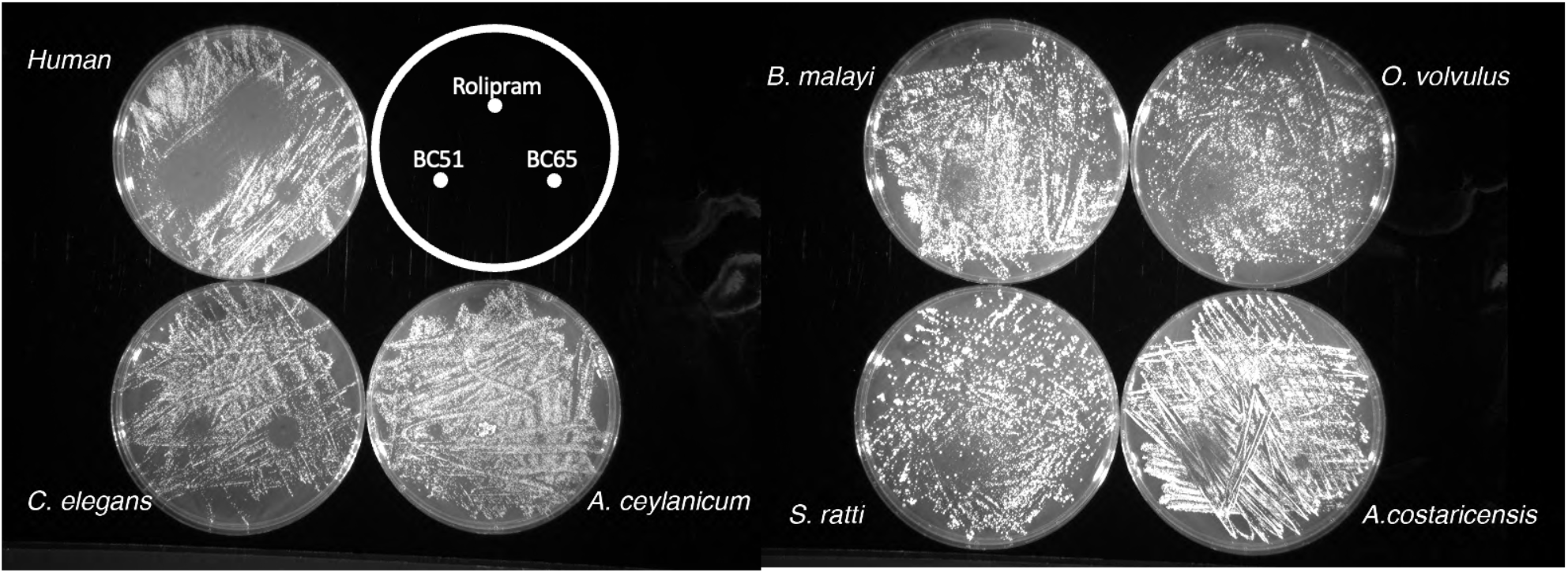
Rolipram, BC51, and BC65 were tested against human PDE4B2, *C. elegans* PDE4, *Ancylostoma ceylanicum* (Acey) PDE4, *Brugia malayi* (Bmal) PDE4, *Onchocerca volvulus* (Ovo) PDE4, *Strongyloides ratti* (Strat) PDE4, and *Angiostrongylus costaricensis* (Acos) PDE4. Rolipram fails to inhibit any of the nematode PDE4s, while BC51 is active against all of the PDE4s.

### 3.2 5FOA Assay Growth Test Results

The 5FOA assay is the opposite of SC-ura halo assays as it detects compounds that lead to growth. This test is required as halo assay tests require PDE inhibition and diffusion while it is known that there are good inhibitors that fail to diffuse in the halo assays.

First, we carried out a 5FOA assay on strains expressing either the human PDE4B2 or the *C. elegans* PDE4 (Figure 6). Consistent with the halo assay results, Rolipram is highly effective on the human PDE4B2, but shows no activity on the two nematode PDE4s. Meanwhile, BC51 appears to be toxic at high concentrations, but then can be seen to promote growth at lower concentrations in strains expressing human or *C. elegans* PDE4. Furthermore, BC75 and the PAN PDE compound show significant activity against the *C. elegans* PDE4 (Figure 6).

**Figure 6:**
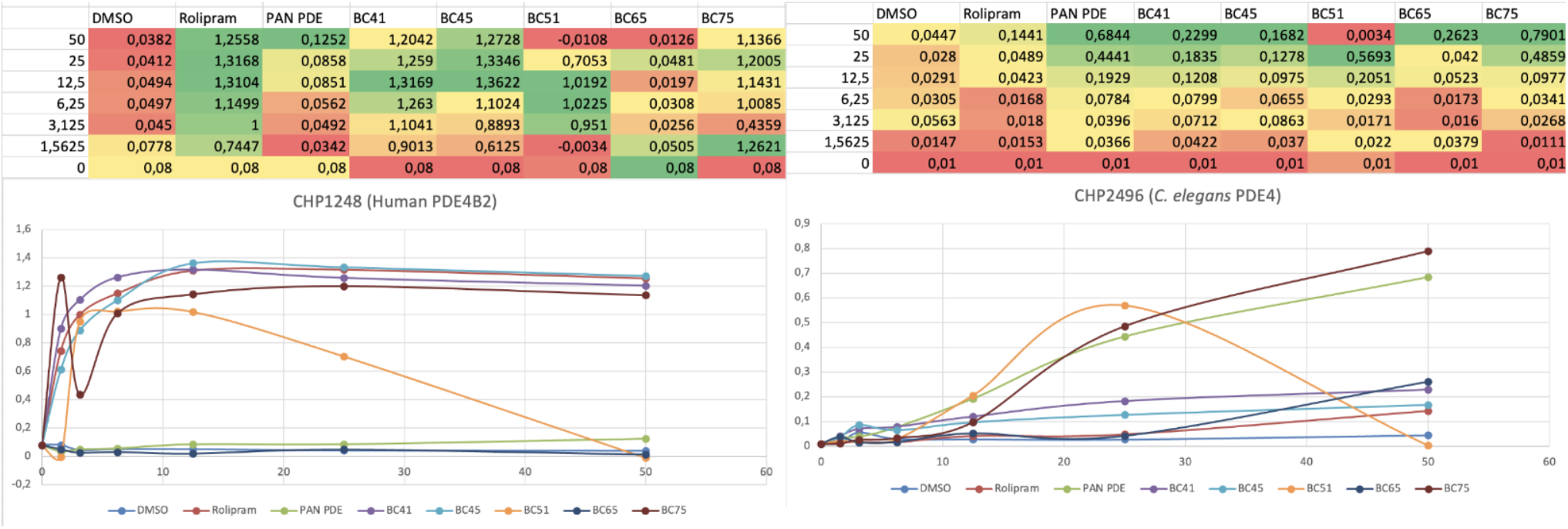
5FOA assay test on strains expressing human PDE4B2 and *C. elegans* PDE4. Rolipram does not confer growth to strain expressing *C. elegans* PDE4. The low value for BC51 at 50μM is due to toxicity.

Both the human PDE4B2 and the *C. elegans* PDE4 are very active enzymes, allowing us to use standard 5FOA conditions. As many of the parasitic nematode PDE4s appeared to be less active, we carried out 5FOA assays in which we adjusted the conditions to optimize PDE inhibitor detection (Appendix A.4). The two parameters we adjusted were the amount of 5FOA and cell density (Figures 7-9). However, not a lot of growth does not mean the assay is not working. We perhaps do not have the particular compounds for the PDE.

**Figure 7:**
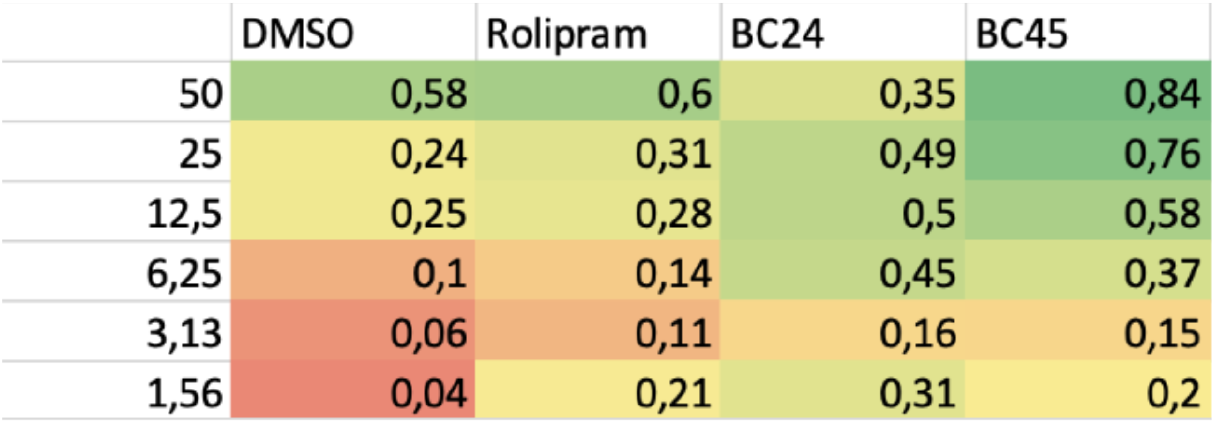
Table shown is a result of 5FOA assay test on a strain expressing *Strongyloides ratti* (Strat) PDE4 using 0.6 g/L 5FOA medium and 2.5 × 10^5^ cells/ml starting density. Rolipram shows no activity, while compounds BC24 and BC45 appear to be active, although BC24 shows toxicity at the highest concentration.

**Figure 8:**
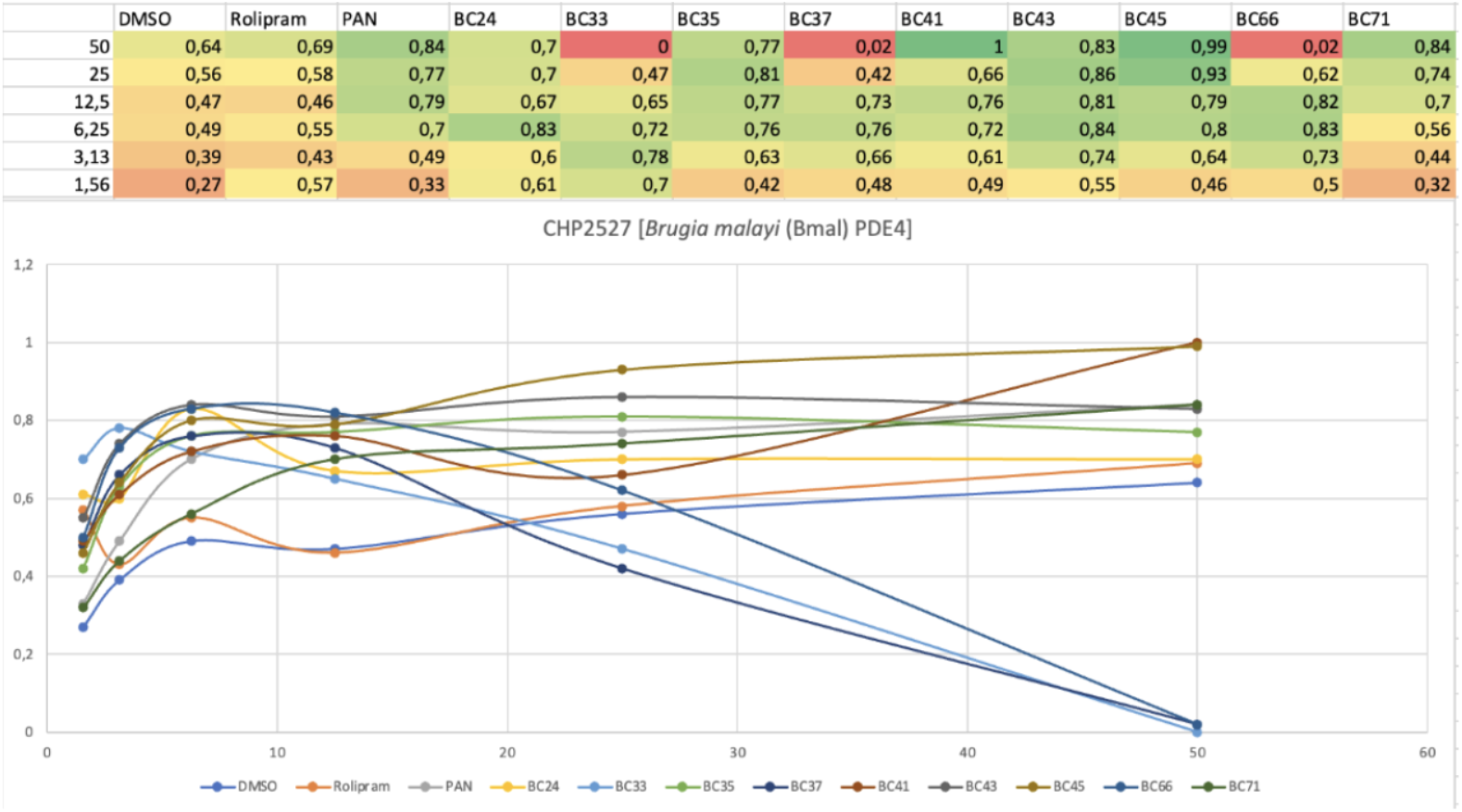
5FOA assay test on *Brugia malayi* (Bmal) PDE4 using 0.55 5FOA medium and 2.5 × 10^4^ cell density. All of the compounds, except for Rolipram, stimulate growth.

**Figure 9:**
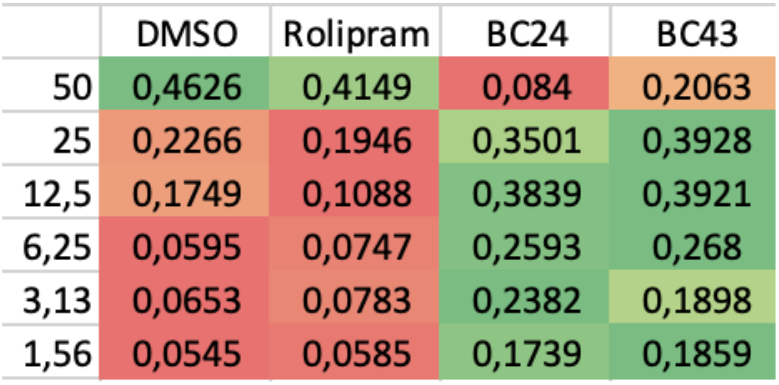
5FOA assay test on a strain expressing *Onchocerca volvulus* PDE4 using 0.6 g/L 5FOA medium and 2.5 × 10^4^ cells/ml starting density. Rolipram shows no activity, while compounds BC24 and BC43 appear to be active, although both show toxicity at the highest concentration.

## 4 Discussion

Our hypothesis of this project was that compounds designed against mammalian PDE4s do not necessarily work on nematode PDE4 and our goal of this project was to identify compounds that do work on nematode PDE4s.

During this research, we demonstrated that nematode PDE4s do not show the same activity as mammalian PDE4 inhibitors. Assays involving very active PDEs require low levels of 5FOA, but permit high starting cell densities. Assays with strains whose PDEs appear to be less active require higher levels of 5FOA to inhibit growth in the absence of a PDE inhibitor and may require lower starting cell densities.

Both the Halo Assays and the 5FOA growth assays showed that Rolipram, a drug that defined the mammalian PDE4 family, does not inhibit the nematode PDE4s. This is consistent with the biochemical studies by Schuster et al. (2019) [8] that found that the PDE4 inhibitors roflumilast and zardaverine were approximately 100-fold less effective on *C. elegans* PDE4. Upon aligning the sequences of the catalytic domains of human PDE4B2 and the six nematode PDEs used in this study (Appendix A.5), one sees that amino acid 407 (threonine) of the human enzyme is different from those in the nematode PDEs. A mutation in PDE4B2 that changed this threonine to an alanine was shown to confer resistance to Rolipram [14], thus explaining why so many PDE4 inhibitors fail to act on the nematode PDE4s.

In this research, we were able to identify compounds that worked on each of the nematode PDE4s. In the 40 compound halo assay (Figure 4), BC65 produces halos on all of the strains, although it does not produce halos on the strains expressing the parasitic nematode PDE4s (Figure 5). In the 5FOA assays, various compounds showed the greatest activity against individual PDE4s. For example, BC45 is one of the best inhibitors of PDE4s from *Strongyloides ratti* and *Brugia malayi* (Figures 7 and 8), while BC75 is one of our most effective inhibitors of the *C. elegans* PDE4. These results suggest that we could evaluate the effect of PDE4 inhibition of the relevant nematodes using these compounds.

In parasitic nematode (Figure 8), BC43 and BC45 seem to be the most effective compounds against Bmal PDE4. Meanwhile, BC33, BC37, and BC66 seem to be less effective as it is seen to be toxic at high concentration. And in Strat PDE4 (Figure 7), BC45 is seen to be more effective than BC24.

BC51 worked best on all of the strains in the halo assay. However, the 5FOA assay revealed that it is toxic at high concentrations, while showing PDE inhibition at lower concentrations (Figure 6). Therefore, the halos were actually a combination of general toxicity closer to the compound spot together with PDE inhibition further from the compound. As such, some of the smaller halos might only reflect toxicity. More 5FOA assays will be required to verify BC51 as a PDE inhibitor of all of the nematode PDE4s.

## 5 Future Work

In the future work, in terms of more optimization of 5FOA condition, we can adjust the amount of 5FOA and cells to make it more optimal; the more 5FOA we add, the more we shift to inhibiting growth. The more cells we add, the more we will be able to see growth. The fewer cells we have, the more we shift to a lower density. The less 5FOA we have, the more we allow growth. The optimization of the assay is very strain-dependent [15], with all these different strains (Appendix A.1) we will have different growth as we have different conditions.

Once optimized, 5FOA assays should be carried out using the full set of PDE inhibitors listed in Appendix A.2.

## 6 Conclusion

In this project, we demonstrated successful expression of phosphodiesterases from all the strains (Appendix A.1) and showed that the nematode PDE4s are resistant to most, but not all, mammalian PDE4 inhibitors. This includes Rolipram, the drug used to define the mammalian PDE4 family. The inability to inhibit these PDE4s is likely due to a single amino acid difference at position 407 in the human PDE4B2, which is a threonine in the human enzyme, but an asparagine or a valine in the nematode enzymes (Appendix A5). However, the 5FOA assays did identify compounds from the Hoffman collection that could inhibit various nematode PDE4s, which could be used to evaluate the effects of PDE4 inhibition on *C. elegans* or as a positive control compound in a new high-throughput screen (HTS) for even more potent inhibitors that do not inhibit mammalian PDE4s.

## 7 Practical Takeaways

- Nematode PDE4s are generally insensitive to inhibitors of mammalian PDE4s.
- The fission yeast system can be used to express phosphodiesterases from all the strains (Appendix A.1) and identify compounds that inhibit their activity.
- Compound Rolipram fails to inhibit any of the nematode PDE4s.
- In halo assay, compound BC51 shows activity against mammalian and nematode PDE4s while in 5FOA assay, BC51 shows toxicity at high concentrations and PDE4 inhibition at lower concentrations.
- Nematode PDE4 inhibitors identified in this study can be used to evaluate the effects of PDE4 inhibition on *C. elegans* or as a positive control compound in a new high-throughput screen (HTS) for even more potent inhibitors that do not inhibit mammalian PDE4s.

## 8 Acknowledgments

I would like to express my profound gratitude to my mentor Professor Charles S. Hoffman, Ph. D. of Boston College for his supervision, guidance, and advice throughout my project. I am also grateful for Natalie Chen, Judy Ly, Hannah Sutoris, Maira Ahmed, Jeremy Eberhard, and Nicholas Rhodes of Boston College for giving me a step-by-step guidance in the lab and experiment. I’d like to thank to Mr. Mark Kantrowitz, Ms. Maite Ballestro, Mrs. D, Yoland Gao, Steven Liu and Siya Goel, Barış Ekim, and Donald Liveoak for their invaluable help throughout RSI. I would also like to thank the Massachusetts Institute of Technology (MIT), the Center for Excellence in Education (CEE), and Mr. Nicholas Nash and Ms. Phalgun Raju of Nick Nash and Phalgun Raju Foundation Scholar for providing me with this research opportunity and their sponsors. I’d also like to extend my appreciation to Eli Meyers, *Ayah, Bunda, Mas* Rafi, dr. Dona Arlinda, and *SMA Negeri* 78 Jakarta Teachers. Lastly, I’d like to thank Lintang Alnafoura and Naya Siregar for their continuing support at all times.

## A Appendix

### A.1 Strains name list

| Strain | h | fbp1::ura4 | ura4::fbp1<br>-lacZ | leu1-32 | Additional markers | cAMP pathway<br>mutations |
| --- | --- | --- | --- | --- | --- | --- |
| CHP1268 | - | fbp1::ura4 | ura4::fbp1<br>-lacZ | leu1-32 | pap1D::ura4-cgs2::PDE4B2 | git3::kan |
| CHP2496 |  | fbp1::ura4 | ura4::fbp1<br>-lacZ | leu1-32 | pap1D::ura4- cgs2-2<br>lys2-97::pJV1<br>-Cepde-4d:3 | gpa2::his3+ |
| CHP2517 |  | fbp1::ura4 | ura4::fbp1<br>-lacZ | leu1-32 | pap1D::ura4- cgs2-2<br>lys2-97::pJV1<br>-Aceypde4 | git3::kan<br>gpa2? |
| CHP2532 | + | fbp1::ura4 | ura4::fbp1<br>-lacZ | leu1-32 | pap1D::ura4- cgs2-2<br>lys2-97::pJV1<br>-Acospde4 his3-D1 | gpa2::his3+ |
| CHP2528 |  | fbp1::ura4 | ura4::fbp1<br>-lacZ | leu1-32 | pap1D::ura4- cgs2-2<br>lys2-97::pJV1<br>-Ovopde4 his3? | git3::kan<br>gpa2::his3+ |
| CHP2529 |  | fbp1::ura4 | ura4::fbp1<br>-lacZ | leu1-32 | pap1D::ura4- cgs2-2<br>lys2-97::pJV1<br>-Sratpde4 his3? | git3::kan<br>gpa2::his3+ |
| CHP2527 | + | fbp1::ura4 | ura4::fbp1<br>-lacZ | leu1-32 | pap1D::ura4- cgs2-2<br>lys2-97::pJV1<br>-Bmalpde4 his3-D1 | gpa2::his3+ |

### A.2 Compounds Used in the Experiments

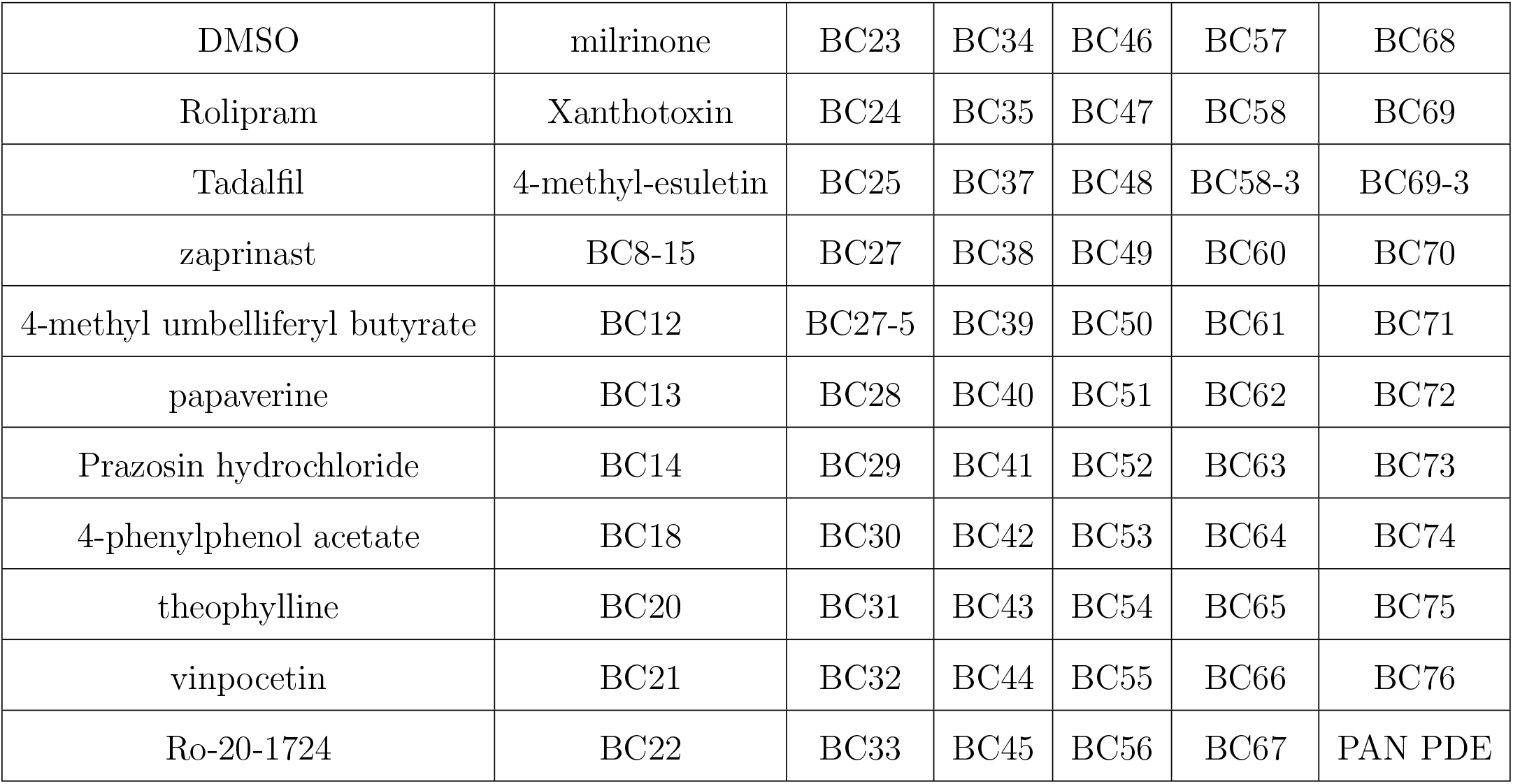

|  |  |  |  |  |  |  |
| --- | --- | --- | --- | --- | --- | --- |
| DMSO | milrinone | BC23 | BC34 | BC46 | BC57 | BC68 |
| Rolipram | Xanthotoxin | BC24 | BC35 | BC47 | BC58 | BC69 |
| Tadalafil | 4-methyl-esuletin | BC25 | BC37 | BC48 | BC58-3 | BC69-3 |
| zaprinast | BC8-15 | BC27 | BC38 | BC49 | BC60 | BC70 |
| 4-methyl umbelliferyl butyrate | BC12 | BC27-5 | BC39 | BC50 | BC61 | BC71 |
| papaverine | BC13 | BC28 | BC40 | BC51 | BC62 | BC72 |
| Prazosin hydrochloride | BC14 | BC29 | BC41 | BC52 | BC63 | BC73 |
| 4-phenylphenol acetate | BC18 | BC30 | BC42 | BC53 | BC64 | BC74 |
| theophylline | BC20 | BC31 | BC43 | BC54 | BC65 | BC75 |
| vinpocetin | BC21 | BC32 | BC44 | BC55 | BC66 | BC76 |
| Ro-20-1724 | BC22 | BC33 | BC45 | BC56 | BC67 | PAN PDE |

### A.3 40 Evaluated Compounds Used in Halo Assays

| Plate 1 |  |  |
| --- | --- | --- |
| Rolipram | BC12 | BC51 |
| BC12 | BC53 | BC14 |
| BC68 | BC15 | BC18 |

| Plate 2 |  |  |
| --- | --- | --- |
|  | BC54 |  |
| BC23 |  | BC37 |
| BC39 |  | BC44 |
|  | BC55 |  |

| Plate 3 |  |  |  |
| --- | --- | --- | --- |
| Ro-20-1724 | BC11 | BC22 | BC24 |
| BC25 | BC46 | BC27 | BC58 |
| BC61 | BC28 | BC63 | BC29 |
| BC34 | BC73 | BC50 | BC75 |

| Plate 4 |  |  |
| --- | --- | --- |
| BC20 | BC52 | BC57 |
| BC65 | BC66 | BC69 |
| BC69-3 | PAN PDE | Tadafil |

### A.4 5FOA condition that were being adjusted

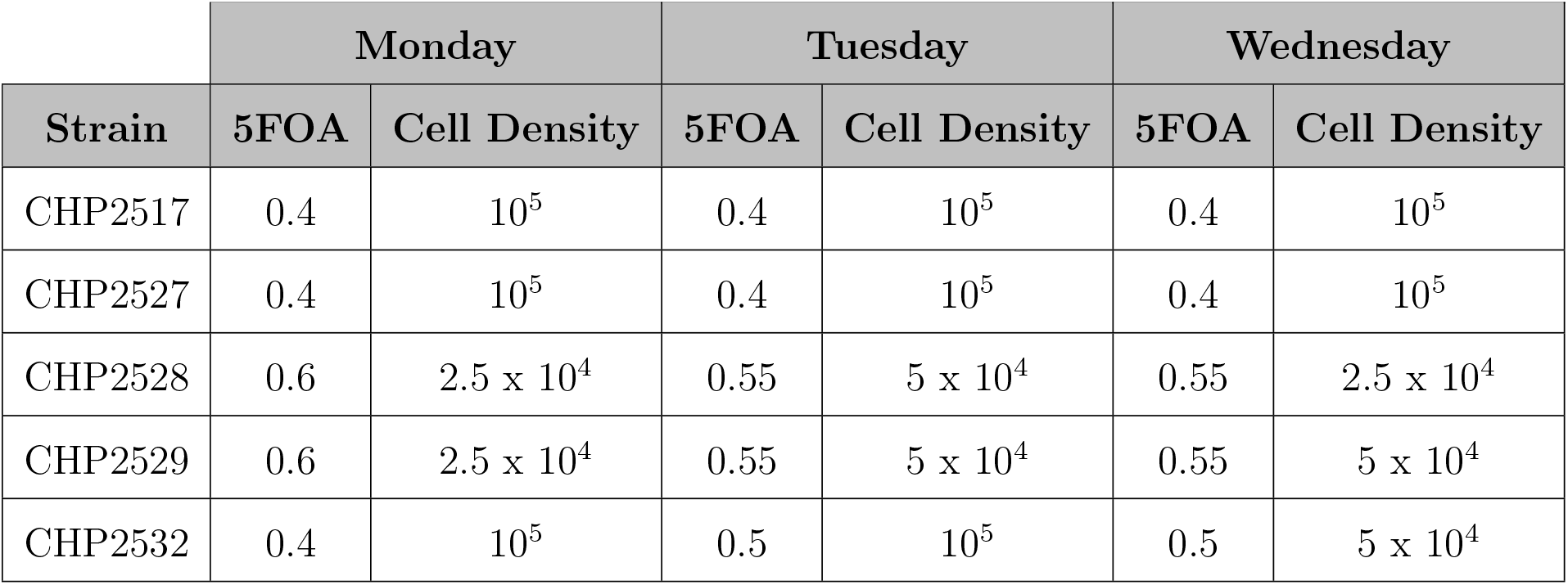

|  | Monday |  | Tuesday |  | Wednesday |  |
| --- | --- | --- | --- | --- | --- | --- |
| Strain | 5FOA | Cell Density | 5FOA | Cell Density | 5FOA | Cell Density |
| CHP2517 | 0.4 | $10^5$ | 0.4 | $10^5$ | 0.4 | $10^5$ |
| CHP2527 | 0.4 | $10^5$ | 0.4 | $10^5$ | 0.4 | $10^5$ |
| CHP2528 | 0.6 | $2.5 \times 10^4$ | 0.55 | $5 \times 10^4$ | 0.55 | $2.5 \times 10^4$ |
| CHP2529 | 0.6 | $2.5 \times 10^4$ | 0.55 | $5 \times 10^4$ | 0.55 | $5 \times 10^4$ |
| CHP2532 | 0.4 | $10^5$ | 0.5 | $10^5$ | 0.5 | $5 \times 10^4$ |

### A.5 Alignment of PDE4 Catalytic Domains from Humans and Nematodes

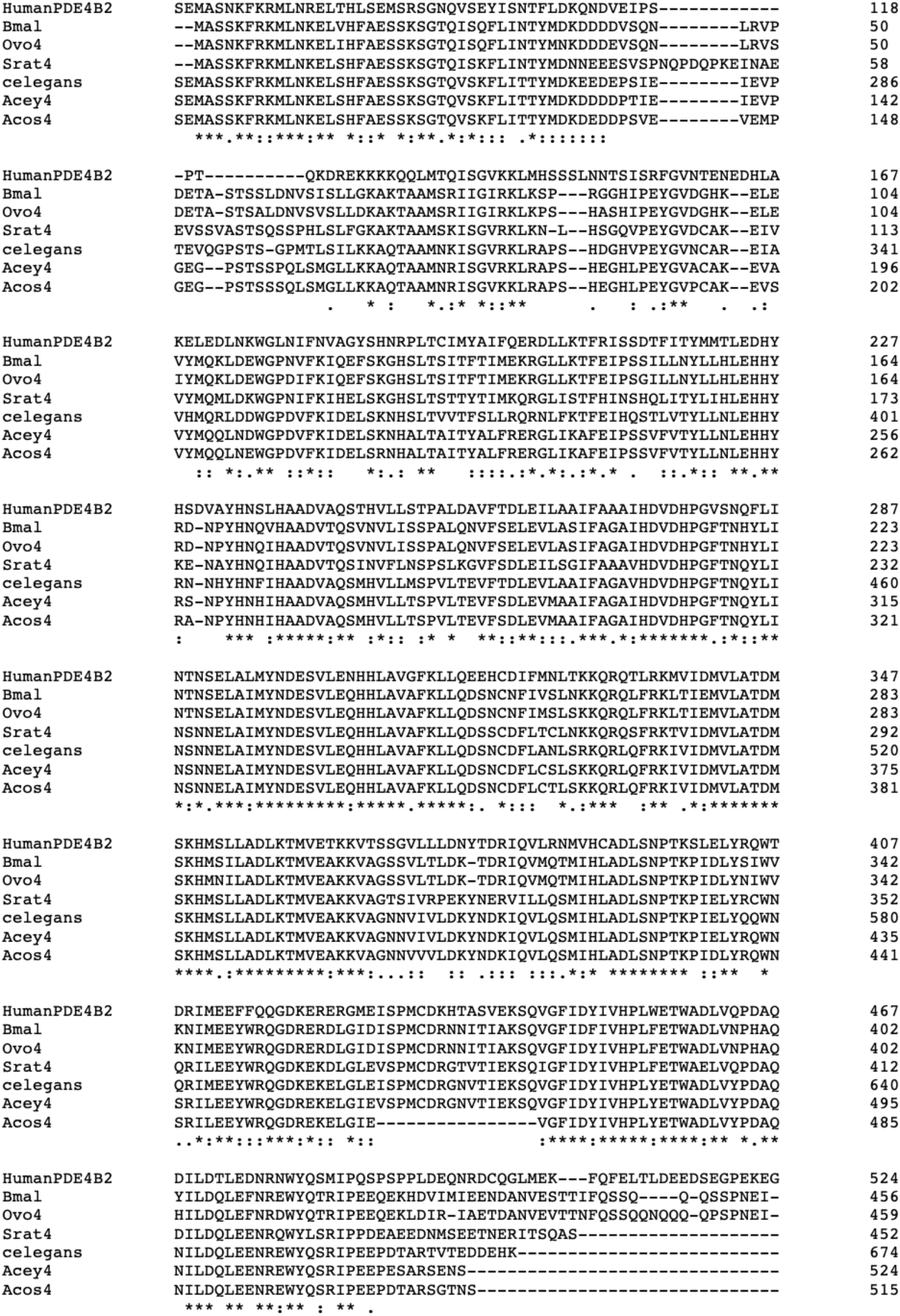

